# Study of Sterilization Effects on Marine *Vibrio* sp. using Interaction of Cavitation with Shock Wave in a Narrow Water Chamber

**DOI:** 10.1101/370445

**Authors:** J. Wang, A. Abe, T. Koita, M. Sun, Y. Wang, C. Huang

## Abstract

When underwater shock waves are generated by an electric discharge in a narrow water chamber, the instant release of a great amount of energy causes the propagation of a shear wave in wall material with the deformation of the chamber wall. The shear waves produce decompression in water and result in the growth of bubble nuclei. Subsequently, those oscillating cavitation bubbles are exposed to the shock pressures, and thus free radicals and rebound shock waves are generated due to their violent collapses. Eventually, marine bacteria around them are inactivated by these productions. In the present study, we investigate the sterilization effects of these oscillating bubbles and cavitation-shock interaction by bio-experiments, respectively. Furthermore, the chemical action of free radicals on marine bacteria is discussed. The generation of the OH radicals is clarified by measuring the concentration of the H_2_O_2_. To estimate the generation condition of the OH radicals, a bubble dynamic model consisting of an oscillation model for the growth of bubble nuclei and an impact model to describe the cavitation-shock interaction is developed. Finally, the theoretical estimation by the bubble dynamic model is discussed under the conditions of the present experiments.

## 1. Introduction

The phenomenon of cavitation was first discovered in 1894 when the tests were made to determine why a ship could not reach its design speed during sea trials. It was found that the generated cavitation makes the hydrodynamic performance of a propeller reduce, while also give rise to vibration and erosion. After that, cavitation bubbles have been observed in many different fields, and their dynamic behaviors have been studied in detail experimentally, theoretically, and numerically. In experiment, there are some generation methods of cavitation bubbles, such as hydrodynamic method including hydraulic and ultrasonic cavitation, and liquid breakdown induced by laser, electric discharge, or underwater explosion. When local static pressure in liquid decreases rapidly below a limiting pressure, the hydrodynamic cavitation occurs and finally would develop into cloud cavitating flow, such as the flows of hydraulic machinery of pump, screws, and water turbines. The collapse and shedding of the cavitation would bring destructive damage to the machinery (1-4). However, the destruction also has a positive effect with appropriate controlling techniques, in the fields of medical therapy, drug delivery, food engineering, waste water treatment, and so on. Takayama (5) found the liquid jet induced by the motion of cavitation bubble near body stone could enhance the effect of extracorporeal shock wave lithotripsy. These cavitation bubbles were generated in tensile region behind underwater shock wave focusing. For drug delivery, the controlled motion of cavitation bubbles by acoustics were proven to enhance drug activity and uptake when high intensity focused ultrasound was introduced to develop a novel technique of targeting drugs to tumors (6). Song et al. (7) applied a high-power laser to the cleaning of the solid surface in liquid and pointed out that a high cleaning efficiency for the removal of particles was obtained by using liquid jets and rebound shock waves induced during the bubble collapse. Loske et al. (8) developed a non-thermal food preservation method using the cavitation-shock interaction and clarified the bactericidal effect of *Escherichia coli* in an electrohydraulic shock wave generator. And they indicated that the bacteria were inactivated by the mechanical action of the shock wave. In the field of maritime sciences, the collapse of microbubble was used to the sterilization of ships’ ballast water (9). When they carried out a bio-experiment to investigate the sterilization effect of the shock wave-microbubble interaction, Wang and Abe (10) found the cavitation bubbles generated behind the concentration of underwater shock waves have a potential to inactivate marine bacteria.

On the other research, Koita et al. (11) observed the generation of cavitation bubbles behind the propagation of multiple waves produced by underwater electric discharge in a narrow water chamber. Regarding the reason of the cavitation generation, they argued that the interactions of the waves between the outer and inner interface of the acrylic wall of the chamber caused decompression in water. To clarify the generation mechanism of these cavitation bubbles, Wang et al. (12) carried out an experiment by producing an air layer between a bag made by silicone film and the water surface in the narrow water chamber in order to remove the action of underwater shock waves, as shown in Fig. 1. The figure shows that a weak transmitted shock wave (TSW) is captured at 30 μs although the propagation of an underwater shock wave is intercepted well by the air layer. Subsequently, a 2nd wave is propagating in the upper side of the air layer and cavitation bubbles are generated behind the 2nd wave. They argued that the 2nd wave was not induced by the permeation of underwater shock wave. On the other hand, as shown in Fig. 1, the elastic wave (ELW) propagating in the wall material is in front of the shock wave (SW) so that the 2nd wave is also not from the reflection of the ELW. They concluded that the 2nd wave causing the generation of cavitation bubbles was a shear wave travelling in the window material. The wave was induced by the deformation of the wall material when large amounts of energy were released as a result of underwater electric discharge. In addition, they also found that interaction of cavitation bubbles with shock waves could enhance the inactivation effect on marine bacteria. Cavitation bubble could be also called as inertial bubble because of its oscillation in the absence of external pressure loading (13-15). Klaseboer et al. (16) investigated the dynamic interaction of a pressure pulse with laser-induced inertial bubble experimentally and numerically, respectively. Their results clarified that the more intense collapses of the bubbles occurred due to the action of shock pressure.

**Fig. 1.**
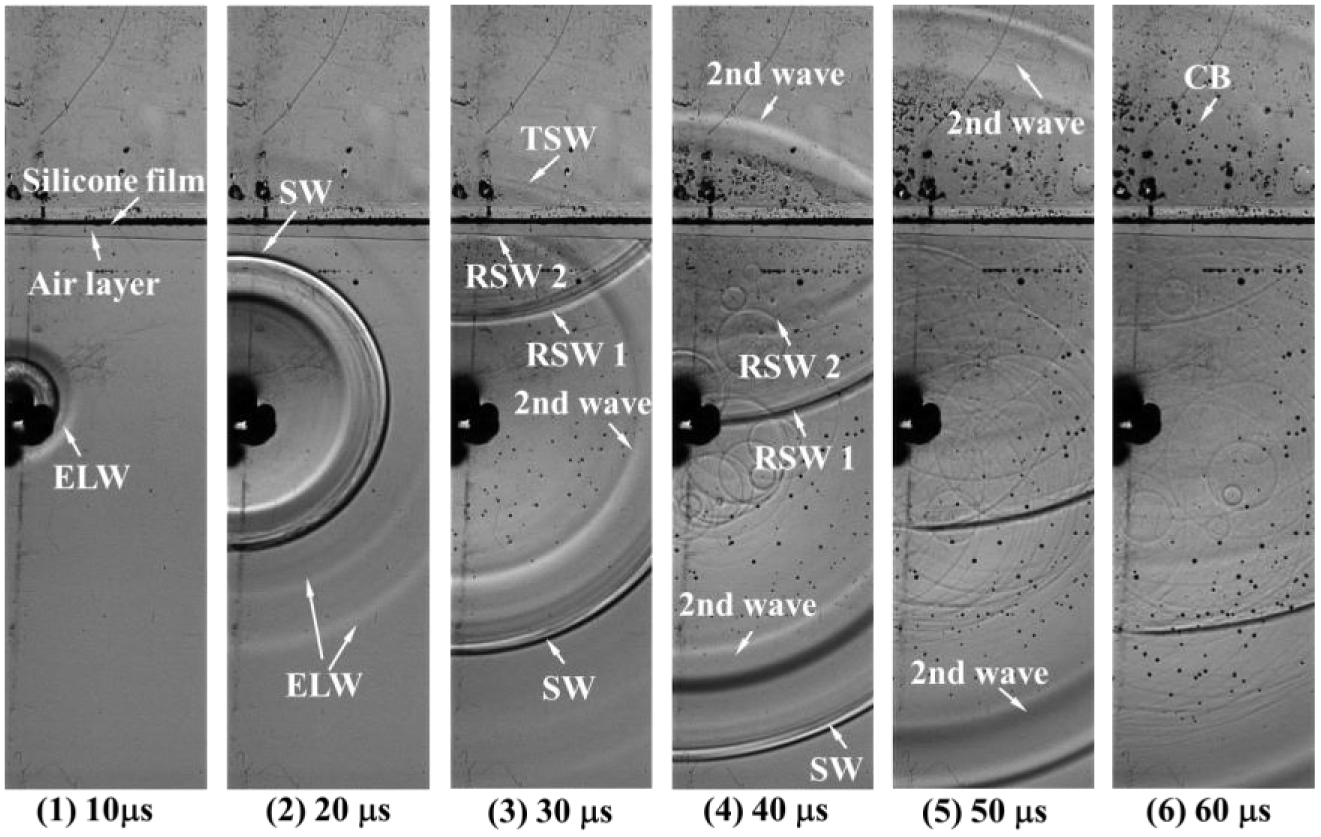
Observation of multiple waves generated by underwater electric discharge with 2-mm air layer in Ref. (12). ELW: Elastic wave, SW: Shock wave, RSW1: Reflection of shock wave, RSW: Reflection of 2nd wave, TSW: Transition of shock wave through air layer, and CB: Cavitation bubbles.

From those backgrounds, considering the sterilizing potential of cavitation bubbles, it is interesting to understand the inactivation effect of the cavitation-shock interaction on bacteria in a narrow water chamber. In bio-experiments of marine *Vibrio* sp., underwater shock waves are produced by a high-voltage power supply, and the cell suspension is isolated from distilled water in the chamber by using a silicone bag. In the experiments, optical observation and pressure measurement are simultaneously conducted to analyze the propagation of underwater shock waves and behaviors of cavitation bubbles. To investigate the sterilization effects of only these oscillating bubbles, an air layer is set to prevent underwater shock waves directly passing through the cell suspension. The respective effects of the chemical and mechanical action of the bubble collapses are also examined. Furthermore, the generation of the OH radicals is clarified by measuring the concentration of the H_2_O_2_. On the other hand, a bubble dynamic model consisting of an oscillation model and an impact model is developed to investigate the condition for generating the OH radicals. Finally, we discuss theoretical solution of the bubble dynamic model under the condition of the present experiments.

## 2. Bio-experimental setup

Figure 2 shows a schematic of the bio-experimental setup in a narrow water chamber. The dimensions of the narrow water chamber were 300 mm (H) × 240 mm (W) × 5 mm (D). Given the observation by Koita et al. (11), it was found that the area at which cavitation bubbles were generated was an annular at a position of about 20 mm from the discharge point. To examine the sterilization effects of these cavitation bubbles, a bag made of a 0.1-mm silicone film was designed in the water chamber and filled with cell suspension of marine *Vibrio* sp., as shown in Fig. 2. The dimensions of the silicone bag were 120 mm (H) × 100 mm (W) × 5 mm (D). Its acoustic impedance is almost the same as that of water. The discharge point was set up at a distance of about 23 mm from the bottom of the silicone bag in the water chamber. Underwater shock waves were continuously generated from the discharge point by a high-voltage pulse power supply (HPS 18K-A, Tamaoki Electronics Co-Ltd) and a pulse generator. In the experiments, the water chamber was filled with distilled water to make sure the triggers of the electric discharges. Its applied frequency was 1 Hz. In addition, optical visualization of schlieren method and pressure measurements were conducted to analyze the propagation behaviors of shock wave and collapsing motion of cavitation bubbles in the water chamber.

**Fig. 2.**
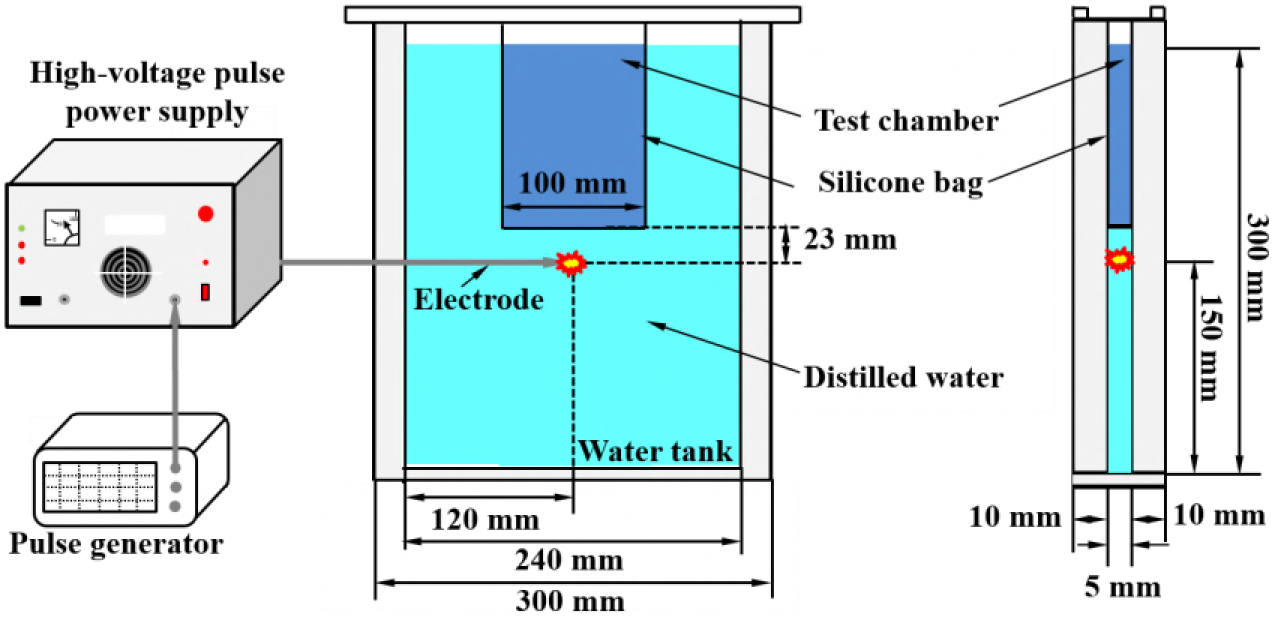
Schematic of bio-experimental setup for estimating sterilization effects

Figure 3 illustrates a considerable sterilization mechanism in the narrow water chamber. Cavitation bubbles are generated behind a shear wave propagating the wall material due to instant release of large amount energy at the discharge point. After the underwater shock waves from the reflection on the chamber wall or the transmission of the elastic wave pass through these oscillating bubble, the intense collapses are induced with the generation of rebound shock waves and free radicals that inactivate marine bacteria by the mechanical and chemical action, respectively. A photo of marine *Vibrio* sp. used in the bio-experiments is shown in Fig. 3 (c). For estimating the sterilization effects of the cavitation-shock wave interaction, cell experiments were carried out in the silicone bag. Samples were extracted regularly from the cell suspension, diluted serially, and spread on the agar plate. The agar places were incubated for 24 hours at 35°C. The cell viability in 1 ml was evaluated using the number of colony-forming cell in the agar places on the basis of the dilution ratio.

**Fig. 3.**
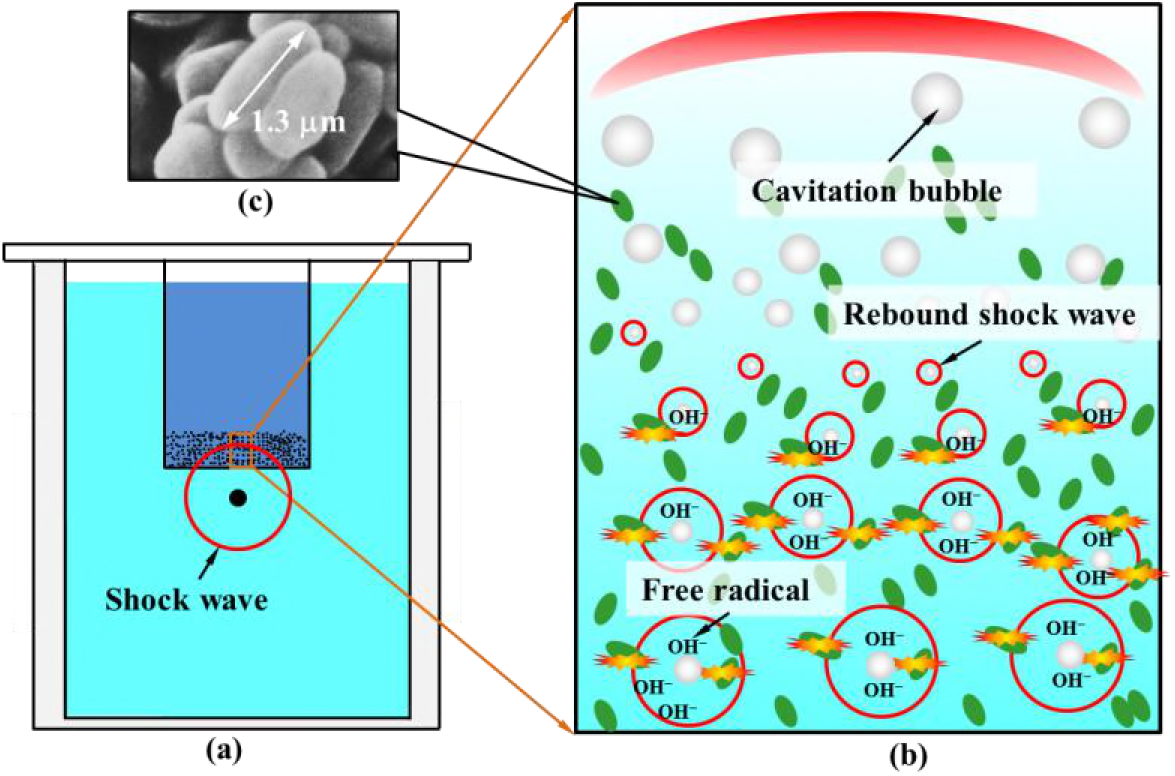
Schematic of inactivation mechanism of cavitation-shock interaction generated in narrow water chamber: (a) Schematic of narrow water chamber, (b) Collapses of cavitation bubbles, a nd (c) Photo of marine *Vibrio* sp.

## 3. Bubble dynamic model

According to our previous study (17), the action of the OH radicals is mainly responsible for inactivating marine bacteria in a cylindrical water chamber at a 31.6-kV electric discharge. Comparing with the experimental conditions in Ref. (17), the shock pressures inducing bubble collapse in the narrow water chamber are obviously lower, especially in the case of setting an air layer, as shown in Fig. 1. Nevertheless, we could obtain a high sterilization effect in Ref. [12], which is thought to depend on the biochemical action of the OH radicals. Consequently, a bubble dynamic model was developed for investigating the condition for the generation of the OH radicals in the interaction of cavitation bubbles and shock pressures, as shown in Fig. 4. From the given generation mechanism, these cavitation bubbles are generated from bubble nuclei when local static pressure is decompressed below the saturated vapor pressure by the tensile action in water with the propagation of a shear wave in the wall material. After that, these bubbles are exposed to the pressure oscillations by reflected shock waves in water. To analyze those collapsing motion, it requires to build an oscillation model for the growth of bubble nuclei and an impact model for the cavitation-shock interaction in the bubble dynamic model. In the oscillation model, the growth of bubble nuclei is thought to be isothermal since the heat transfer is fast relative to the time scale of the bubble motion (18). In the meantime, an equilibrium evaporation and condensation are also considered to investigate the transportation of water vapor at the bubble interface. For the cavitation-shock interaction, it is necessary to analyze the bubble motion after the bubble in arbitrary motion phase is exposed to a shock pressure, as shown in the figure. Here, the heat transfer through a thermal boundary layer is needed to estimate the conditions for generating the OH radicals besides the transportation of water vapor. In the bubble dynamic model, the bubble is assumed to maintain a spherical shape during its collapsing motion.

**Fig. 4.**
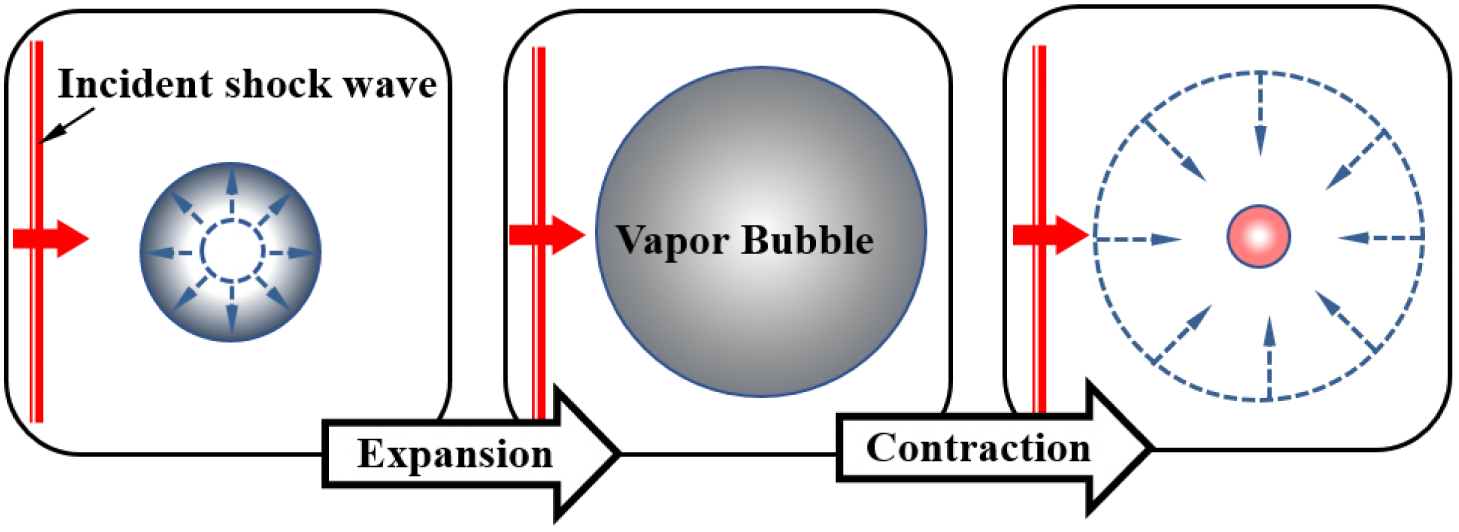
Schematic of bubble dynamic model for describing growth of bubble nuclei and cavitation-shock interaction

### 3.1 Oscillation model

In the model, the Rayleigh-Plesset equation was solved to describe the growth of bubble nuclei, as presented in Eq. (1).

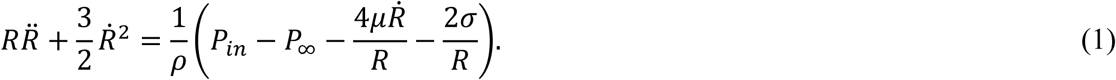

where *R* is the bubble radius, *Ṙ* = d*R*/d*t*, *t* is the time, *Ȑ* = d*Ṙ*/dt, *P*_in_ is the pressure inside the bubble, *P*_∞_ is the pressure behind an incident shock wave, *σ* is the surface tension, and *μ* is the viscosity coefficient.

For the equation for state inside the bubble, the van der Waals equation was used,

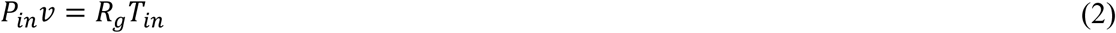

where *v* and *T_i_*_n_ are the molar volume and temperature of the gas inside the bubble, and *R_g_* = 8.3145 J/(mol K). The molar volume *v* is presented by,

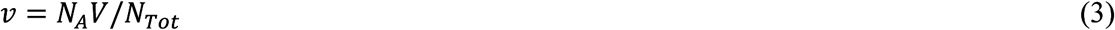

where *N*_A_ is the Avogadro number, 6.02×10^23^, the volume of the bubble *V* = 4/3π*R*^3^, and *N*_Tot_ is the total number of gas molecules in the bubble, consisting of the number of water vapor *N*_water_ and other molecules *N*_others_.

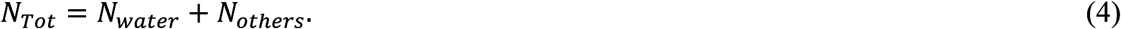

In the model, we just consider the transfer of water vapor through a boundary layer during the growth of a nucleus. Given the study of Toegel et al. (19), the rate of particle change for water vapor can be estimated by Eq. (5).

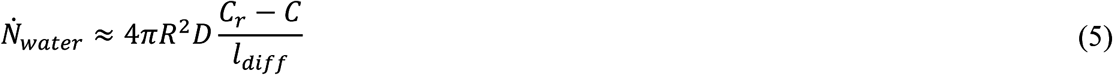

where *C*_r_=*C*_r_ (*T*_0_) corresponds to the equilibrium density of water vapor at the bubble wall (20, 21), *C*_r_ = *P*_v_(*T*_0_)/*kT*_0_ ≈ 5.9×10^23^ m^-3^ for *T*_0_=293.15K, *k* is the constant, 1.38×10^−23^ (22), *C* is the actual concentration of water vapor inside the bubble, *C* = *N*_water_/*V*, *D* is the diffusion constant, and *l*_diff_ is the diffusive penetration depth. In the study of Toegel et al. (22), the diffusive penetration depth can be presented referring to the expression of a thermal boundary layer. Hence, *l*_diff_ could be obtained with the Rayleigh-Plesset time *τ*_c_ (17, 23).

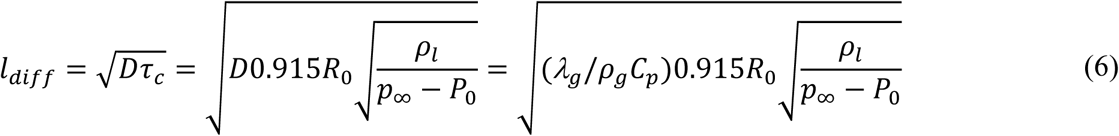

where the diffusive constant *D* is approximately equals to the thermal diffusivity α_g_ = *λg*/*C*_p_*ρ_g_*, *λg* is the thermal conductivity, *C*_p_ the heat capacity at constant pressure, *P*_0_ is the atmospheric pressure, *ρ*_g_ is the density of gas and varies with the temperature and pressure inside the bubble, as shown in Eq. (7).

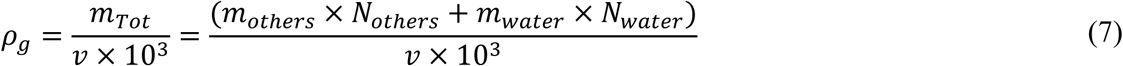

where *m* denotes the molar mass in unit of g/mol.

### 3.2 Impact model

In the impact model, the Herring bubble equation was applied considering the compressible of water when the speed of sound was assumed to be constant (24).

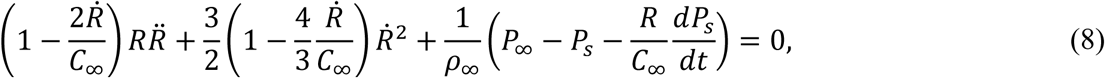

where *C*_∞_ is the sound speed of water at infinity, *ρ*_∞_ is the density of water at infinity, and *P*_s_ is the pressure at the wall of a bubble.

*C_∞_* is given by

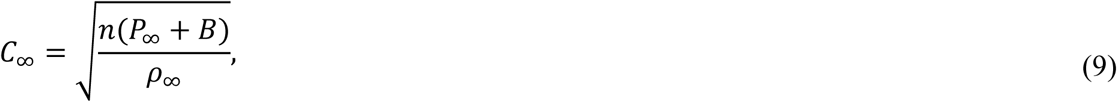

where *B* and *n* are constant values, *B* = 2963 bar and *n* = 7.41.

*P_s_* is described by

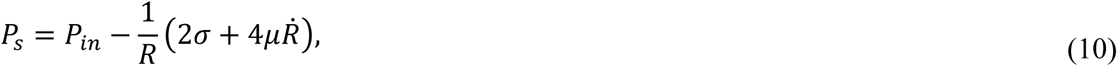

where *P*_in_ is the pressure of the gas inside the bubble.

Besides the transportation of water vapor through the diffusive penetration depth, as expressed in Eqs (4)-(7), the effect of the thermal conduction at the bubble wall was also required in the impact model. Hence, the energy balance of the gas inside the bubble is written in Eq. (11) according to the first law of the thermodynamics, where the bubble radius varies from *R* to *R*+Δ*R* and the gas temperature changes by Δ*T* during the time Δ*t*:

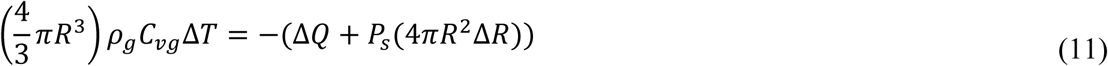

where *C*_vg_ is the specific heat of gas at constant volume. The term on the left-hand side presents the variation of the internal energy of the gas during the time Δ*t*, the second term on the right-hand side is the work of the pressure force at the bubble surface, and Δ*Q* is the heat released from the bubble to the liquid through a thermal boundary layer. The boundary layer is basically formed in the gas inside the bubble since the density and specific heat of water are so much larger than the respective values for gas. In the model, it is assumed that the temperature is spatially uniform at the center of the bubble, while to be linear within the boundary layer. Hence, Δ*Q* can be written in Eq. (12) according to the Fourier’s law,

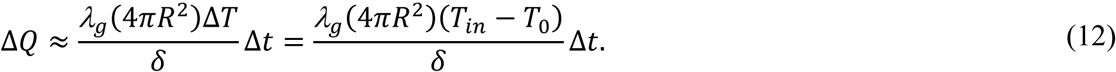

where *δ* is the thickness of the thermal boundary layer, *δ* ≈ *l*_diff_, as mentioned above. *T*_0_ and *T*_in_ are the temperature of the gas inside the bubble at *t* = 0 and *t* + Δ*t*.

Finally, the Rayleigh-Plesset and Herring bubble equation were solved using the fourth-order accurate Runge-kutta-Gill method

## 4. Results and discussion

Figure 5 shows multiple waves generated by an underwater electric discharge in the narrow water chamber, observed using the schlieren method. The output power of the electric discharge was 28.6 kV. The optical schlieren method was carried out using a metal halide lamp (LS-M350, SUMITA optical glass Inc.). The high-speed camera (i-SPEED 7, Nac Image Technology) captured images at a frame rate of 100 kfps and an exposure time of 300 ns. The resolution of the images was 840 × 216 pixels. In the figure, a 1st shock wave (SW) generated by the electric discharge is observed in Fig. 5 (1). The 1st SW is thought to be cylindrical due to the thickness of the water chamber. The images indicate that its propagation speed is about 1500 m/s. At 20.87 μs, the 1st SW is reflected partly at the bottom of the silicone bag, as indicated by Reflection 1, and then transmitted through the silicone film. After a wave passes through the film, an obscure outline of its reflected wave (Reflection 2) is also observed. According to our previous observation (12), the wave is thought to an elastic shear wave (SHW) generated by the deformation of the wall material due to the instant release of enormous energy when the electric discharge was triggered. From Fig. 5, it can be seen that cavitation bubbles (CB) are generated and grows behind the SHW in the upper side of the silicone film. Subsequently, rebound shock waves (CSW) are captured when the reflected shock waves (RSW) from the both sides of the water chamber pass through these cavitation bubbles, as shown in Fig. 5 (7). These observations indicate the inactivation of marine bacteria is expected in the cavitation-shock interaction. In addition, the pressure measurement of FOPH 2000 (Fiber Optical Probe Hydrophone, RP acoustic) was simultaneously conducted with the observation of the schlieren method.

**Fig. 5.**
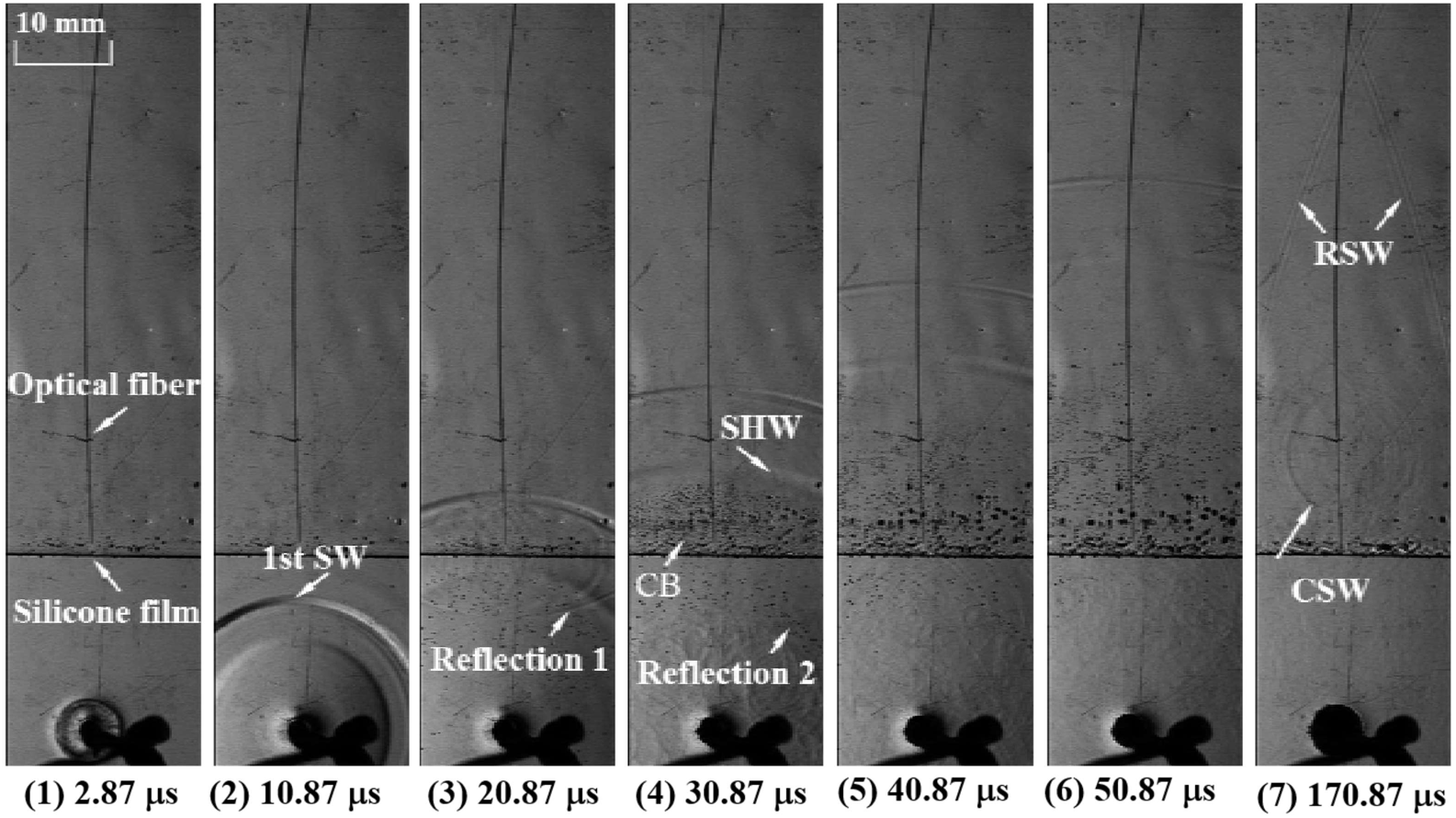
Observation of multiple waves generated by underwater electric discharge in narrow w ater chamber using schlieren method

Figure 6 shows a pressure profile obtained at a distance of 23 mm from the discharge point using the FOPH 2000. In the figure, the experimental data within 2.5 μs are affected by the flash noise of the electric discharge. We observed the 1st SW of about 9.5 MPa at 15.4 μs and the SHW of about −7.4 MPa at 24.4 μs. As indicated by the red arrows, there are some pressure fluctuations within the area between the 1st SW and SHW. According to the observation in Fig. 5 (3), it is thought that they show the pressure records of rebound shock waves generated by the collapses of air bubbles attaching to the silicone bag from the beginning when exposed to the 1st SW. The pressure variations after 30 μs are due to the motion of the cavitation bubbles. On the other hand, we should note the accuracy of the negative pressure of the SHW since the pressure data measured by the FOPH 2000 is estimated using the Tait equation. The density *ρ* in the tensile region could be obtained to about 996.2 kg/m^3^ when the atmosphere *P*_0_ = 1.01325×10^5^ Pa, the negative pressure, *P*_n_ = −7.4×10^6^ Pa, and the density at the atmosphere *ρ*_0_ = 999.7 kg/m^3^.

**Fig. 6.**
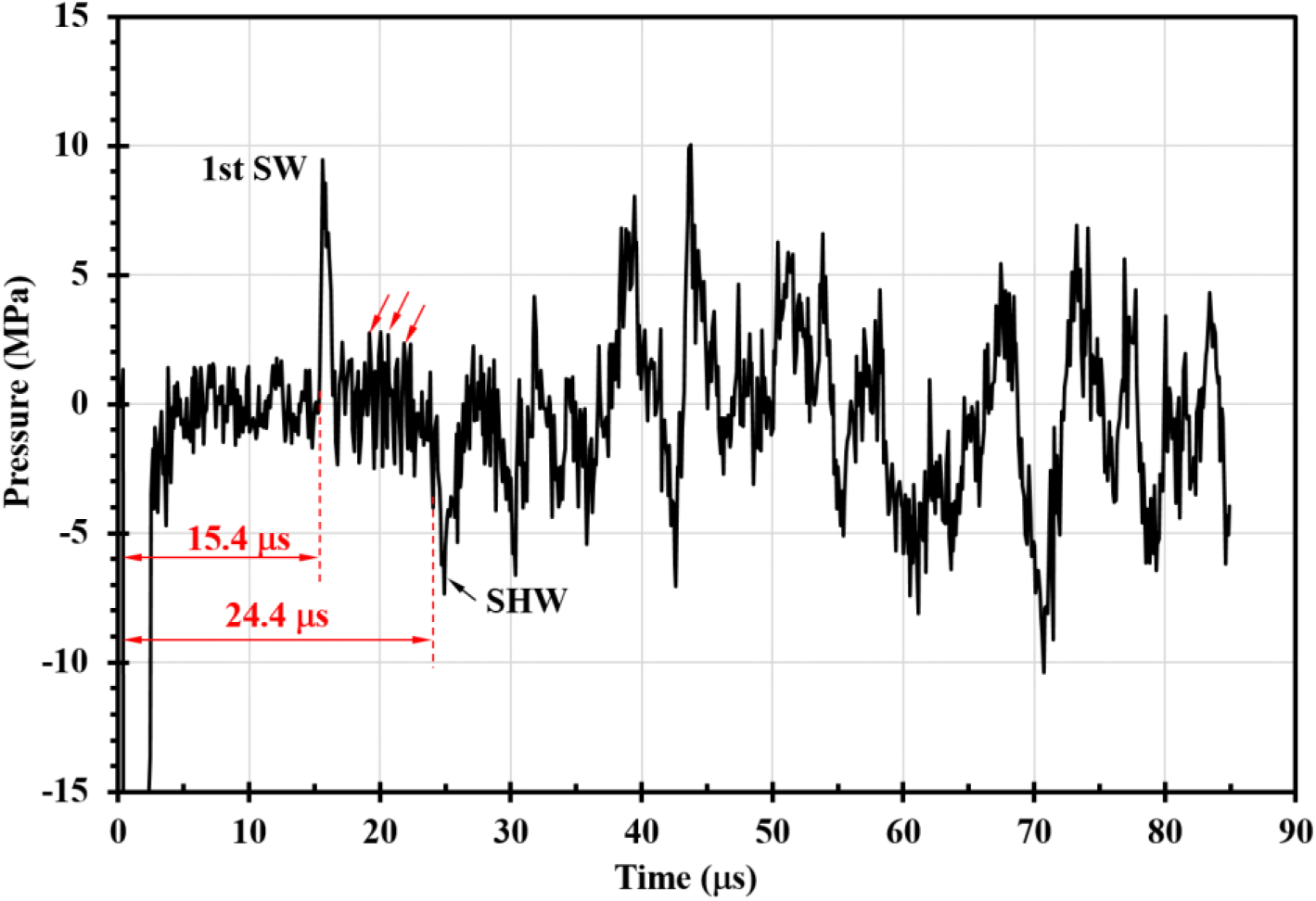
Pressure profile obtained using FOPH 2000 at position of about 23 mm from discharge point

Figure 7 shows estimates of the number of viable cells for an electric discharge of 28.6 kV. The initial concentration of marine bacteria was about 5.37 ×10^4^ cfu/ml. The plots shown in this figure are of the averages for six sets of the bio-experimental data. Here, the error bars are not shown because the values of the standard deviation (STD) are too small to be clearly recognized in the exponential ordinate of the viability ratio. In the figures including the following bio-experimental results, the STD are less than 11, even close to 0 except the point (STD ≈ 22) at the beginning of the experiments. The solid squares are the reference data obtained from the cell suspension without the electric discharges. The number of viable cells hardly changes throughout the experiment. To attain the sterilization effect of only these cavitation bubbles, an air layer of 2 mm was set to prevent underwater shock wave entering the upper side of silicone film, as shown in the figure. The solid triangles and diamonds represent the results of the bio-experiments obtained with and without an air layer, respectively. The results show that after 4 min, all of the marine bacteria are completely inactivated without air layer, while only one order of the number of marine bacteria are inactivated with a 2-mm air layer. It indicates the interaction of the cavitation bubble with the shock pressures enhance the sterilization effects. On the other hand, the solid triangle data indicate a tendency that the inactivation rate is slow but the number of viable cells is definitely reduced just by those oscillating bubbles. Their motions are also captured as shown in Fig. 8. The frame rate of the high-speed camera was 300 kfps. From Fig. 8, it is observed that the bubble motions are induced successively by interaction with the rebound shock waves of other bubbles. As a result, a shock pressure leading to the collapse of the bubble plays an important role in a high sterilization effect.

**Fig. 7.**
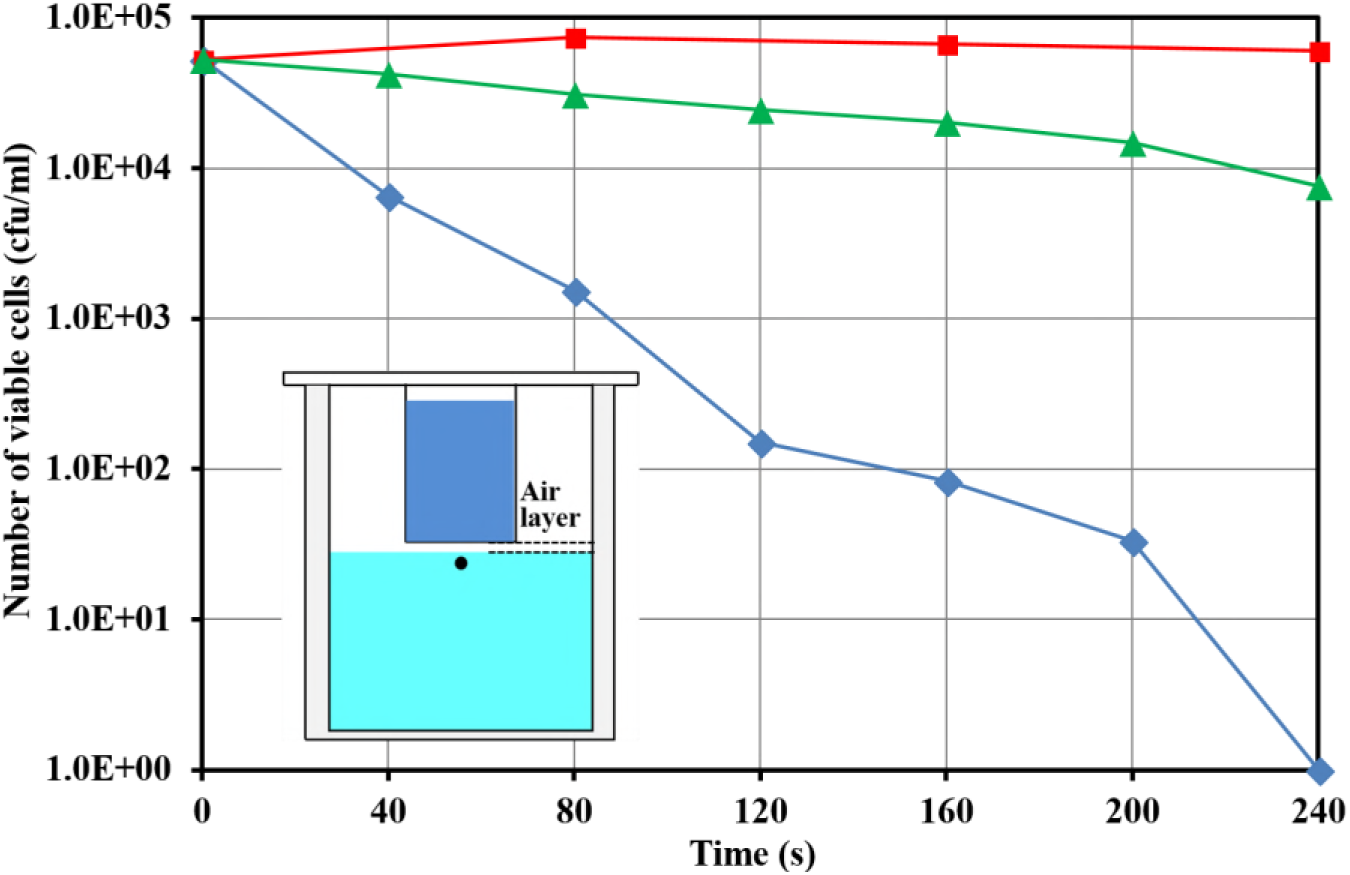
Estimation of sterilization effect obtained with 28.6-kV electric discharges: 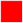 reference data, 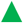 with 2-mm air layer, and 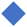 without air layer

**Fig. 8.**
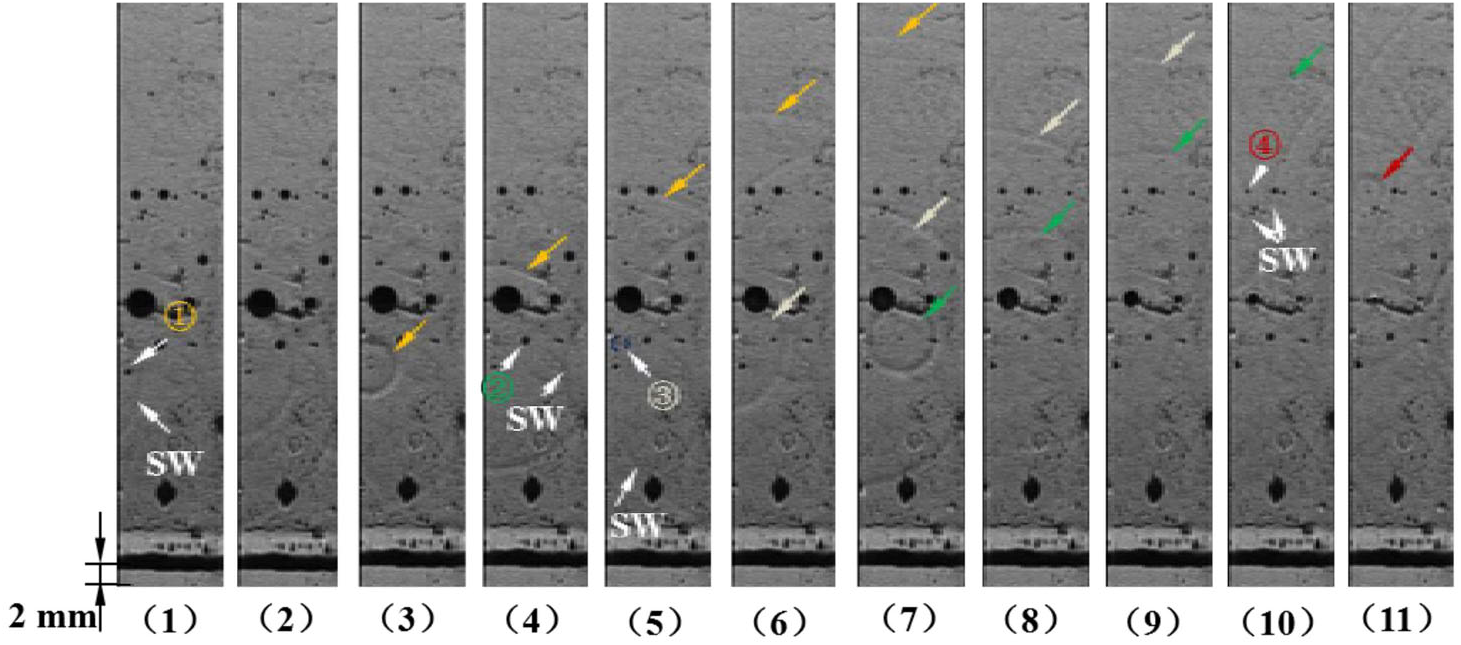
Sequential images of collapses of cavitation bubbles with 2-mm air layer: the interval time is 3.33 μs

Figure 9 shows estimation on the number of viable cells under different output powers of the electric discharges. The solid triangles, diamonds, and circles represent the results obtained with an electric discharge of 31.6 kV, 28.6 kV, and 18.8 kV, respectively. From Fig. 9, it can be seen that the times to obtain perfect inactivation are 160 s for 31.6 kV and 240 s for 28.6 kV, while about two orders of the number of marine bacteria were inactivated after 240 s for 18.8 kV. The sterilization effect obviously increases with the output power of the electric discharge. It goes without saying that a larger output power of the electric discharge makes underwater shock waves stronger. In addition, the instant release of the discharge energy can deform the wall material and spread the width between the walls. Consequently, water in the narrow chamber is instantly stretched with the wall deformation, and a larger number of cavitation bubbles are also generated. On the other hand, as mentioned in Fig. 3, marine bacteria are inactivated by the mechanical action of rebound shock wave and the biochemical action of free radicals produced in the cavitation-shock interaction process. To investigate their respective sterilization effect, the sodium L-ascorbic was put into the cell suspension to remove the effect of free radicals. The results at 31.6-kV electric discharges with the sodium L-ascorbic are presented by the solid squares. These data show that the number of marine bacteria barely changes during the experiment. It indicates that marine bacteria are not inactivated by the mechanical action of the bubble motion, and free radicals take mainly responsible for the inactivation in the present experimental setup. According to our previous study (10, 25), it has been found that the strong rebound shock waves are not generated in the present experimental chamber. In addition, considering the experimental results with an air layer of 2 mm in Fig. 7, all of the marine bacteria are also inactivated by the bio-chemical action of free radicals.

**Fig. 9.**
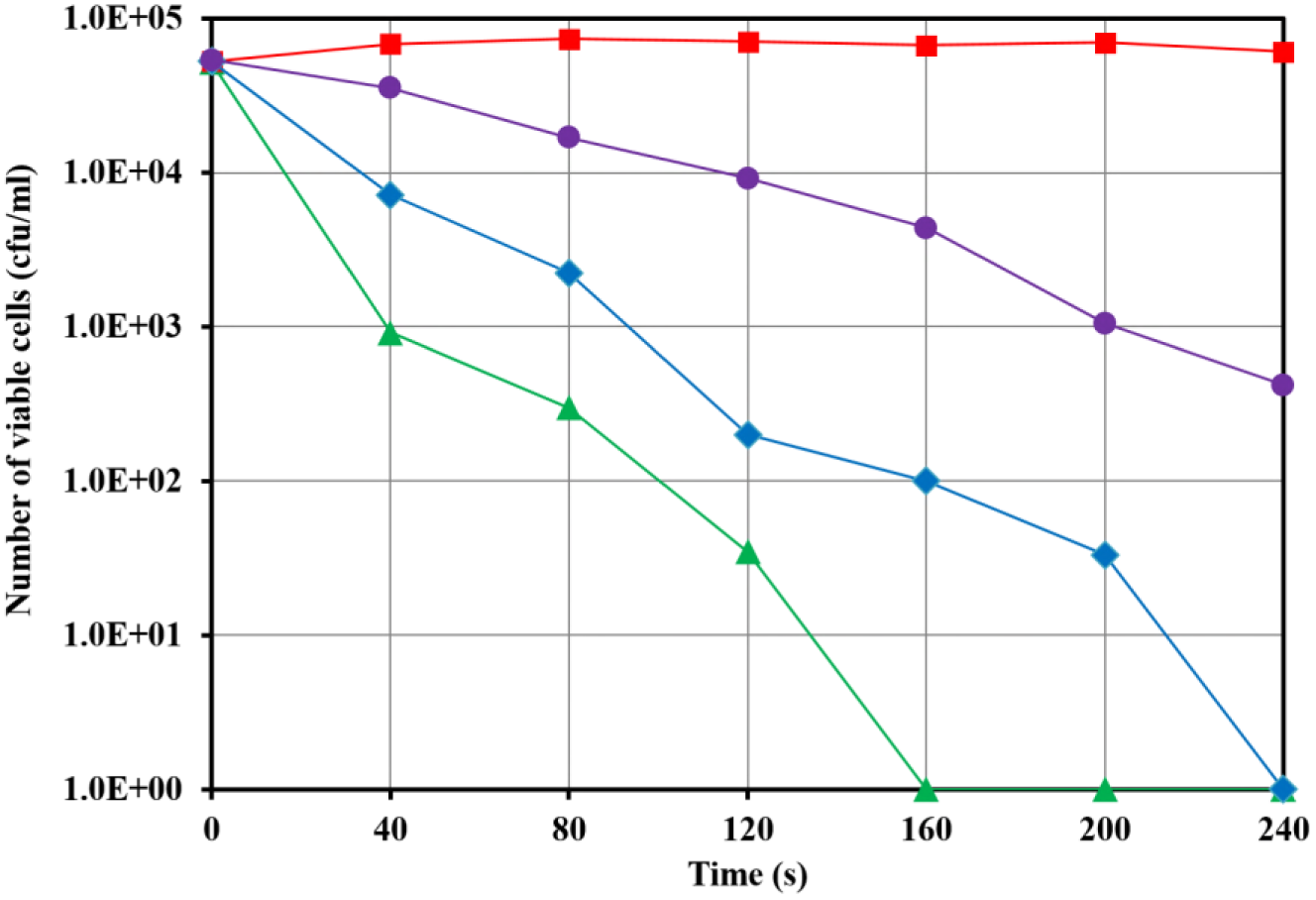
Estimation of number of viable cells obtained with electric discharges of 31.6 kV to 18.8 kV: 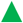 31.6 kV, and 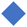 28.6 kV, 18.8 kV, 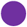 and 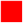 with sodium L-ascorbic

Free radicals inactivating marine bacteria are generated inside the bubble due to extremely high temperature and pressure, and then transfer to the outside through a boundary layer. Among the production of the chemical processes, hydroxyl (OH) radicals are thought to be dominant specie due to their highly oxidative ability. However, it is difficult to directly measure or observe the OH radicals owing to their fast reaction in a dynamic stimulus (26). To examine the generation of the OH radicals in the present study, a digital pack tester (DPM-H_2_O_2_, Kyoritsu Chemical-Check Lab., Corp.) were used to measure the concentration of the H_2_O_2_, one of the productions of their chemical reactions. Its measuring range is from 0.10 to 2.0 mg/ L and the resolution is 0.05 mg/L. Figure 10 shows the concentration of the H2O2 with an electric discharge of 31.6 kV to 18.8 kV. The measurements were conducted after the electric discharges of 5000, 8400, and 10200 shots. The symbols in the figure indicate of the averages for 5-set measurements that are all the same owing to the respectively small resolution of the digital pack tester. The results show that the concentration of H_2_O_2_ increases with the output power of the electric discharge. It indicates an increase in the strength of the cavitation-shock interaction. In the case of a 2-mm air layer, the concentration of the H2O2 is detected to about 0.5 mg/L after 10020-shot electric discharges, so that marine bacteria are inactivated only by those oscillating bubbles, but a high sterilization effect is not obtained. The results of the measurements show good agreements with the bio-experimental results.

**Fig. 10.**
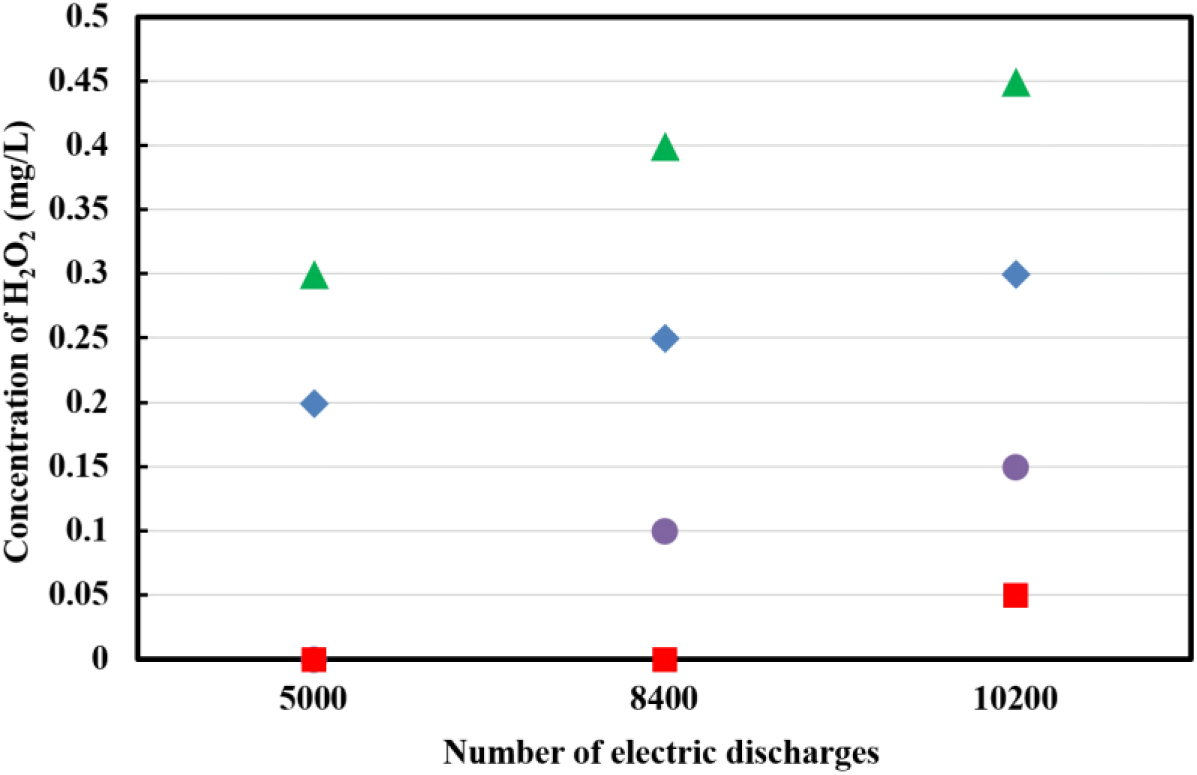
Estimation of concentration of H_2_O_2_ using digital pack tester with electric discharge of 31.6 kV to 18.8 kV: 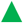 31.6 kV, and 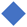 28.6 kV, 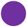 18.8 kV, and 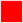 with 2-mm air layer

Based on the pressure measurements, the pressure of reflected shock waves are only dozens of atmospheres, even a few atmospheres in the case of the air layer in the narrow water chamber. However, we obtain a high sterilization effect using the chemical action of the OH radicals in the cavitation-shock interaction, as shown in Fig. 9. To estimate the condition for generation of the OH radicals, a theoretical analysis was carries out by solving the bubble dynamic model. First of all, the theoretical solution of the Rayleigh-Plesset equation for the growth of a 5-μm radius bubble nucleus is shown in Fig. 11. In this analysis, the density of water *ρ*_∞_ = 999.7 kg/m3, the viscosity coefficient *μ* = 1.307 ×10^−3^ Pa·s, the surface tension *σ* = 74 × 10^−3^ N/m, the density of gas *ρ*_g_ = 1.20 kg/m^3^ at *T*_0_ = 293.15 K when *R* = *R*_0_, the thermal conductivity in the gas *λ_g_* = 0.599 W/(m·K), the heat capacity of gas *C*_p_ = 4186 J/(kg·K), the atmospheric pressure *P*_0_ =1.01325 ×10^5^ Pa were used. The initial total number of the molecules inside the bubble was determined by Eqs. (11) and (12) when *P*_in_ = 1.03×10^5^ Pa, *T*_in_ = 293.15 K, *R* = 5 μm. The initial number of water vapor *N*_water_ corresponded to the equilibrium density at the wall, *C*_r_. The difference between the number of total molecules and water vapor was defined as the number of other molecules, *N*_others_. In this figure, the blue lines and orange lines indicate the results obtained with and without the mass transportation, respectively. For the variation of the bubble radius indicated by the solid lines, the bubble nucleus grows to a larger size, about 13 μm in radius in the case of the transportation of water vapor at the interface. Correspondingly, the minimum internal pressure is 0.077×10^5^ Pa, slightly higher than 0.069×10^5^ Pa when not considering the mass transportation due to the energy loss of transferring water vapor, as shown by the dashed lines. Furthermore, the minimum internal pressure slightly increases at the second expansion. Hence, the pressure difference between inside and outside of the bubble shows the largest at the first expansion, so that maximum values of temperature and pressure in the bubble are obtained when the oscillating bubble is exposed to a shock pressure. As a result, the motion phases of the bubble during the first expansion in Fig. 11 were substituted into the impact model in order to analyze the interaction of a bubble with a shock pressure.

**Fig. 11.**
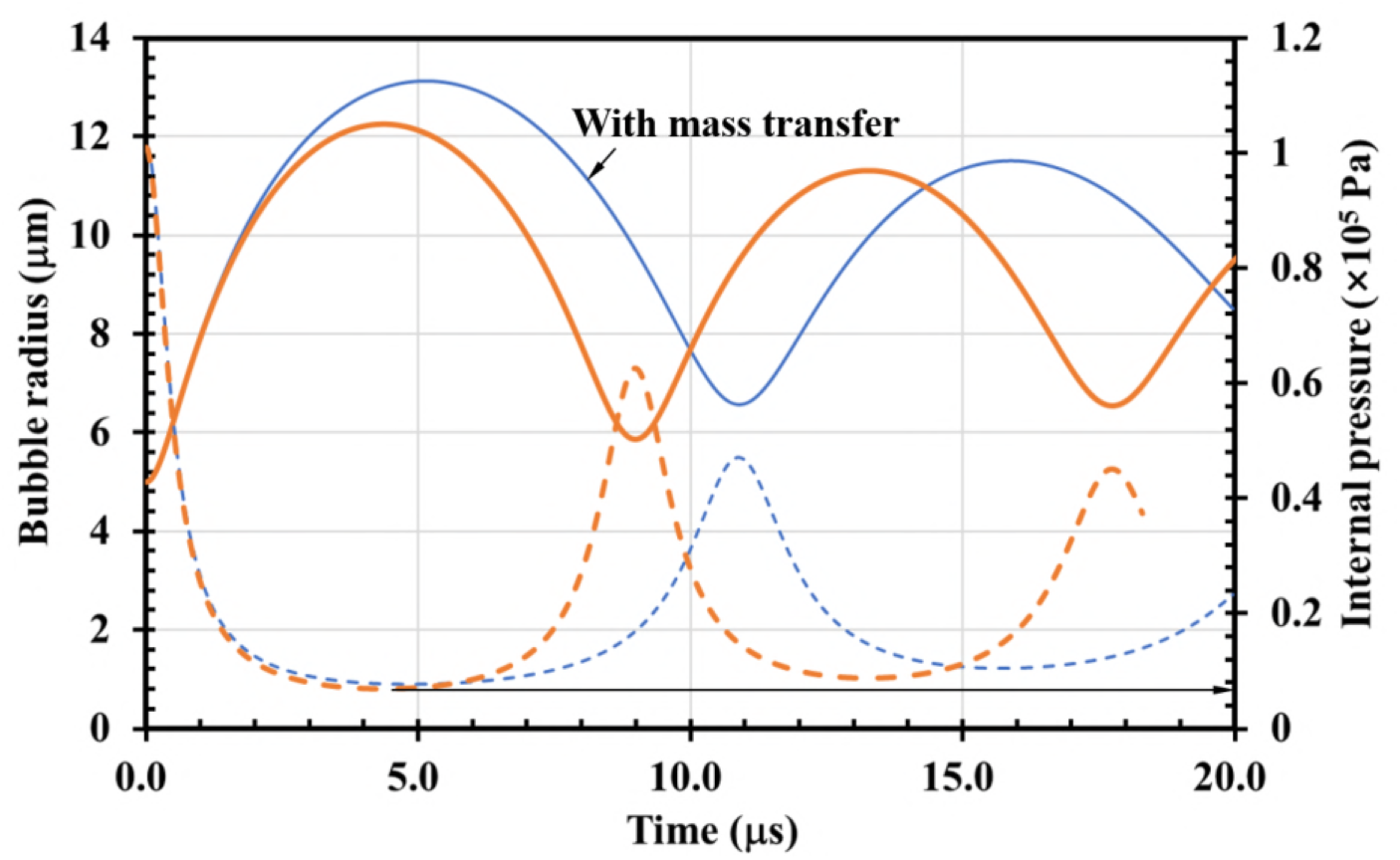
Oscillation of 5-μm radius bubble nucleus solved by Rayleigh-Plesset equation with and without mass transportation

In the present water chamber, there are possibilities that shock pressures interact on oscillating bubbles in arbitrary phases as to radius, surface velocity, and internal pressure. To obtain the collapsing motions of these bubbles, the Herring equation was solved in consideration of heat conductivity and mass transfer in the impact model. The theoretical solution is shown in Fig. 12. The abscissa indicates the time corresponding to the initial bubble motion phases in Fig. 11, and the ordinate is the peak internal temperature of the bubble obtained when the oscillating bubble in this motion phase is exposed to a shock wave of *P*_shock_ = 2 MPa. The pressure of 2-MPa is the average of reflected shock wave at 28.6-kV electric discharge. In the figure, it can be seen that the peak internal temperature is the smallest value of about 1400 K at the starting point; *R*_0_ = 5 μm, *u* = 0 m/s, *T*_in_ = 293.15 K, and *P*_in_ = 1.01×10^5^ Pa. During the expanding phase of the bubble, the peak internal temperature increases due to violent bubble motion induced by a large pressure difference between inside and outside of the bubble, Δ*P_s-i_* = *P*_shock_ − *P*_in_, and reaches a maximum value of about 7000 K at *R*_0_ = 13 μm, *u* ≈ 0 m/s, *T*_in_ = 293.15 K, and *P*in = 0.077×10^5^ Pa. When the shock pressure interacts with a contracting bubble, it is found that the internal peak pressure decreases to about 2400 K at *R*_0_ = 6.59 μm, *u* ≈ 0 m/s, *T*_in_ = 293.15 K, and *P*_in_ =0.48×10^5^ Pa. On the other hand, we should note the conditions of the bubble just when exposed to the shock wave. The radius, surface velocity, and internal pressure of bubble mainly influence the surface tension, 2*σ* /*R*_0_, the kinetic energy, *ρ*_l_ *u*^2^, and the pressure difference between outside and inside of the bubble, Δ*P_s-i_*. Here, the *ρ*_l_ *u*^2^ and 2*σ* /*R*_0_ range within 0 - 0.12 ×10^5^ Pa and 0.11×10^5^ Pa - 0.29×105 Pa during the first expansion of the bubble, as shown in Fig. 11. Compared with the value of the pressure difference, the surface tension and kinetic energy value slightly works on the calculation of the peak internal pressure. As a result, it is obviously concluded that the pressure difference, *ΔP_s-i_*, is a main factor on the collapse of the bubble when exposed to a shock pressure.

**Fig. 12.**
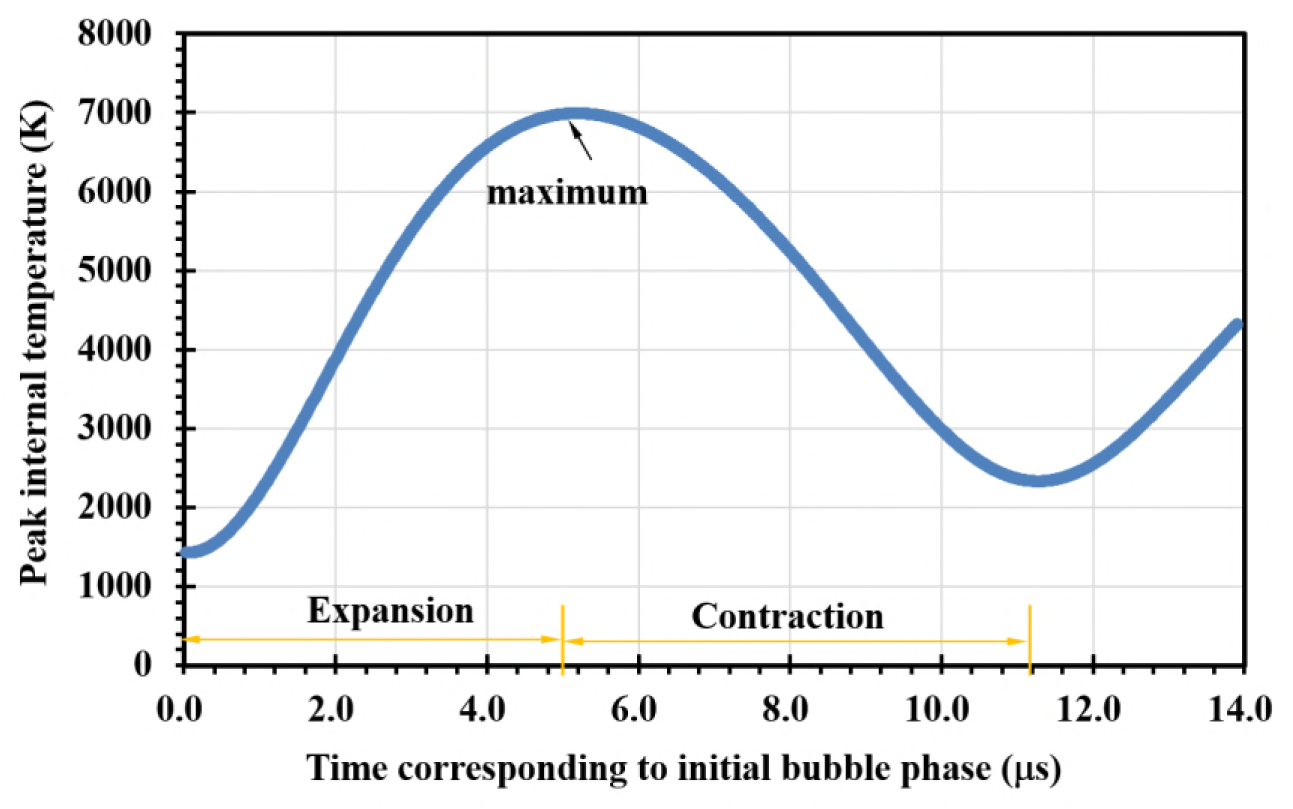
Peak temperatures inside bubbles corresponding to initial states interacting shock pressure of 2 MPa

From the given experimental observation, the pressures leading to the collapses of oscillating bubbles are mainly from the reflected shock waves on the chamber wall or the transmission of the elastic wave travelling in the wall material. The experimental pressure measurements indicated that the pressure ranges of shock waves interacting with oscillating bubbles probably reach about dozens of atmospheres. Hence, we assumed that *P*_shock_ ranges from 0.1 MPa to 5.0 MPa and analyzed theoretically the maximum internal temperatures and pressures generated by the collapse of bubbles exposed to the shock pressures. The results are shown in Fig. 13. The maximum of the peak internal temperatures as indicated by an arrow in Fig. 12 are represented as a maximum internal temperature at *P*_shock_ = 2 MPa in Fig. 13. Corresponding maximum internal pressures are also solved by the bubble dynamic model. From Fig. 13, we can see that the maximum internal temperatures and pressures produced by the cavitation-shock interactions are estimated to be over 1200 K and 50 MPa in the range of *P*_shock_ = 0.1 to 5.0 MPa, respectively. As described in the study of Wang et al. (17), the OH radicals could be generated at the internal temperature of bubble, above 1000 K. Incidentally, the condition of 0.1-MPa is equivalent to the experimental condition of a 2-mm air layer. In this case, underwater shock waves are prevented to entering the upper side over a silicone film. However, cavitation bubbles are still produced due to the shear waves propagating in the wall material. During the growth of the bubble nuclei, it is thought that the ambient pressure around them could return rapidly to atmosphere. Hence, we assumed that the oscillating bubbles are suddenly exposed to a 0.1-MPa ambient pressure. Consequently, as shown in Fig. 10, we obtained the generation of the OH radicals in evidence, but it is difficult to say that we could indicate significant results in the experiment under the condition of an air layer. On the other hand, from Fig.13, the maximum temperature and pressure in a bubble increase with the increase of the shock pressure and reach about 10800 K and 15600 MPa at *P*_shock_= 5 MPa, respectively, so that the concentration of the H_2_O_2_ is probably increased with increase of the inducing shock pressures and leads to a higher sterilization. Those results are supported by the experimental results indicated in Figs. 9 and 10. Hence, the generation of the OH radicals can be obtained at *P*_shock_ = 0.1 MPa, and their concentration increases with increase of the shock pressure in the cavitation-shock interaction. The theoretical solutions obtained with the bubble dynamic model are consistent with the bio-experimental results and concentration measurements.

**Fig. 13.**
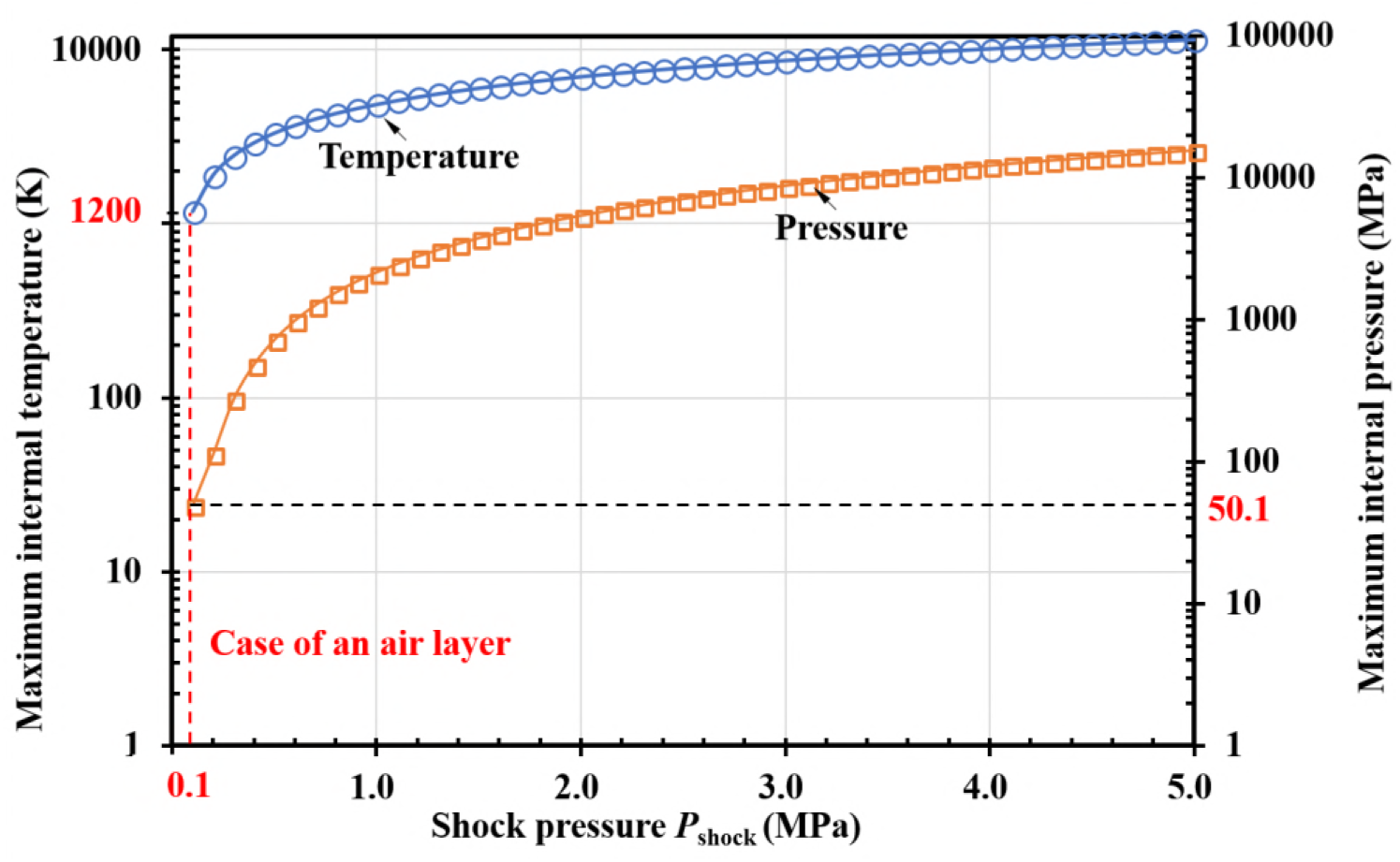
Theoretical solution for interaction of oscillating bubble with shock pressure of 0.1 MPa to 5 MPa

## 4. Summary

Sterilization effects of the cavitation-shock interaction were investigated by a bio-experiment of marine *Vibrio* sp. in the narrow water chamber. Cavitation bubbles were generated by a tensile force acting on water due to the shear waves travelling in the window material as a result of instant release of electric discharge energy. The observation of the schlieren method showed that the collapses of those cavitation bubbles were induced by the underwater shock wave, and their interaction has a potential to inactivate marine bacteria. In the bio-experiment, an air layer was set to prevent the underwater shock waves entering the cell suspension, so that the sterilization effects of only cavitation bubble were obtained. It was found that all of marine bacteria were perfectly inactivated in several minutes in the case of the cavitation-shock interaction while only one order of bacteria was inactivated with an air layer. It indicated that a high sterilization requires a strong shock pressure leading to violent collapses of cavitation bubbles. Furthermore, the sterilization effects increase with the output power of the electric discharge since both of the shock pressures and the number density of cavitation bubbles increase.

On the other hand, the inactivation of marine bacteria was proven to mainly depend on the bio-chemical action of free radicals, by adding sodium L-ascorbic to the cell suspension. To clarify the generation of the OH radicals, we measured the concentration of the H_2_O_2_ under different conditions of the electric discharges. The concentration of the H_2_O_2_ increased with the output powers. It suggested that a strong strength of the cavitation-shock interaction produces a great number of oxidative radicals, and thus leads to a high inactivation of marine bacteria. The measurements showed good agreements with the bio-experiments.

In order to estimate the condition for generating the OH radicals, a model of the bubble dynamics was developed consisting of an oscillation model for the growth of bubble nuclei and an impact model to describe the interaction of a cavitation bubble with a shock pressure. The transfer of heat and water vapor were also considered in the model. The theoretical analysis was carried out under the conditions of the present experiments. The theoretical solutions showed that the pressure difference between inside and outside of a bubble is a main factor in affecting its collapsing motion. It was also found that the internal temperature and pressure could achieve the condition for the generation of the OH radicals in the case of using an air layer and increase with the inducing shock pressure, so that a higher sterilization effect was obtained, identically as described in the bio-experiments and the concentration measurements.

## Author Contributions

J.W. wrote the main text, and made the bio-experiments and calculation of the bubble dynamic model with the help of T.K. and M.S.. A.A. and Y.W. checked and corrected the work. Furthermore, Yiwei Wang has the idea of this work. C.H. helped in the proper organization of the calculations.

## Acknowledgements

This work was supported by the National Natural Science Foundation of China, grant numbers 11772340 and 11332011, and the JSPS KAKENHI, grant Numbers 16H04600 and 16K14512. In addition, we would like to sincerely thank Nac Image Technology Inc. for the supply of optical experimental equipment.

